# Life-stage and sex influence *Philornis* ectoparasitism in a Neotropical woodpecker (*Melanerpes striatus*) with essential male parental care

**DOI:** 10.1101/2021.12.22.473830

**Authors:** Joshua B. Lapergola

**Affiliations:** Department of Neurobiology and Behavior, Cornell University, Ithaca, NY; Bird Population Studies program, Cornell Lab of Ornithology, Ithaca, NY, USA; Department of Ecology and Evolutionary Biology, Princeton University, Princeton, NJ, USA

**Keywords:** Adult birds, Botflies, Caribbean, Dominican Republic, Myiasis, Parasite-host Interaction, Picidae

## Abstract

The nestlings of many Neotropical bird species suffer from *Philornis* (Diptera: Muscidae) ectoparasitism. While nestlings are typically considered the intended targets, recent work indicates that *Philornis* infest adult birds more frequently than previously appreciated, yet few studies have concurrently surveyed nestlings and adults for *Philornis* in the same population. Over six field seasons (2012–2017), I documented the presence of current or recent subcutaneous *Philornis* infestations on adult and nestling Hispaniolan Woodpeckers *Melanerpes striatus* from the same population in the central Dominican Republic. I tested the following three non-mutually exclusive hypotheses regarding occurrence of *Philornis* on adult birds: (1) nestlings are more vulnerable to *Philornis* parasitism than adults, (2) nesting is associated with *Philornis* parasitism in adults, and (3) *Philornis* parasitism is associated with incubation and brooding investment. While nestling and adult woodpeckers exhibited similar prevalence of parasitism, parasitized nestlings hosted on average 3.5 times more *Philornis* wounds (larvae plus empty wounds) than parasitized adults. Nesting *per se* was not significantly associated with parasitism among adults, as breeding and non-breeding adults showed similar prevalence and intensity. However, nests with *Philornis*-infested young were significantly more likely to have one or both parents also be infested in contrast to nests with infestation-free young. Furthermore, adult males, which perform overnight incubation and brooding, were significantly more likely to be parasitized than adult females. This last result supports the hypothesis that incubation and brooding investment increase the risk of *Philornis* parasitism for adults, but this conclusion is complicated by the lack of an association between parasitism and nesting status. Together, these results raise questions about the degree of host life-stage specialization and whether adult parasitism is incidental or part of an alternative parasitic strategy for *Philornis.*

## Resumen

Los pichones de muchas especies de aves Neotropicales sufren de ectoparasitismo por *Philornis* (Diptera: Muscidae). Mientras que los pichones se consideran típicamente los objetivos previstos, trabajos recientes indican que *Philornis* infestan aves adultas con más frecuencia de lo que se pensaba anteriormente, sin embargo, pocos estudios han examinado simultáneamente pichones y adultos para *Philornis* en la misma población. Durante seis temporadas de campo (2012–2017), documenté la presencia de infestaciones subcutáneas recientes o actuales de *Philornis* en adultos y pichones de la misma población del pájaro carpintero *Melanerpes striatus* en el centro de la República Dominicana. Probé las siguientes tres hipótesis no mutuamente excluyentes con respecto a la aparición de *Philornis* en aves adultas: (1) los pichones son más vulnerables al parasitismo de *Philornis* que los adultos, (2) la anidación está asociada con el parasitismo de *Philornis* en adultos, y (3) el parasitismo de *Philornis* es asociado con la incubación y la inversión de crianza. Mientras que los pichones y los adultos exhibieron una prevalencia similar de parasitismo, los pichones parasitados hospedaron en promedio 3.5 veces más heridas de *Philornis* (larvas más heridas vacías) que los adultos parasitados. La nidificación per se no se asoció significativamente con el parasitismo entre los adultos, ya que los adultos reproductores y no reproductores mostraron una prevalencia e intensidad similares. Sin embargo, los nidos con pichones infestadas de *Philornis* tenían significativamente más probabilidades de tener uno o ambos padres también infestados, en contraste con los nidos con pichones libres de infestación. Además, los machos adultos, que realizan incubación y empollando durante la noche, tenían una probabilidad significativamente mayor de ser parasitados que las hembras adultas. Este último resultado apoya la hipótesis de que la inversión de incubación y de empollando aumentan el riesgo de parasitismo por *Philornis* en adultos, pero esta conclusión se complica por la falta de una asociación entre el parasitismo y el estado de anidación. Juntos, estos resultados plantean preguntas sobre el grado de especialización de la etapa de vida del hospedador y si el parasitismo de adultos es incidental o parte de una estrategia parasitaria alternativa para *Philornis*.

## INTRODUCTION

Nestlings of many bird species suffer from myiasis, “the infestation of healthy or necrotic tissue…by dipteran larvae” (Little 2009 p. 546:546), and in the Neotropics, *Philornis* (Diptera: Muscidae) botflies are the primary cause of healthy tissue myiasis (Teixeira 1999, Dudaniec & Kleindorfer 2006). The larvae of at least 23 *Philornis* species are subcutaneous, blood-feeding parasites (Common *et al*. 2019). Botfly effects on nestlings can be severe (reviewed in Dudaniec & Kleindorfer 2006), in some cases reducing survival (Delanoy & Cruz 1991, Rabuffetti & Reboreda 2007, Hayes *et al*. 2019). Indeed, native and introduced *Philornis* have been implicated in the decline of several island endemic birds, most notably in the Galápagos where introduced *P. downsi* have impacted many endemic species (Fessl *et al*. 2006, Kleindorfer & Dudaniec 2016, Leuba *et al*. 2020). Yet the extent of *Philornis* infestation’s ecological impacts remains poorly known, especially in these botflies’ native ranges. Addressing these gaps will be important for not only advancing basic ornithology but also for understanding whether to account for and how to control *Philornis* in conservation and management efforts.

One aspect of *Philornis* parasitism that requires deeper exploration is the degree of host life stage specialization. The prevailing wisdom has been that *Philornis* target altricial and semi-altricial nestlings while the occasional observations of larvae on adult birds represent opportunistic or misdirected infestation (Teixeira 1999). Some researchers have even posited that *Philornis* cannot successfully pupate once host birds have fledged (Arendt 1985a). Understanding the degree to which *Philornis* parasitizes nestlings and adults has important ramifications for bird populations since nestling parasitism directly impacts reproductive success while adult parasitism could impact survival and reproductive success. In a recent review of published records and analysis of new data from adult capture records from three Caribbean islands, Quiroga et al. (2020) reported adult parasitism for 15 bird species representing 12 families and four orders. While these results indicate that adult parasitism by *Philornis* might be more than opportunistic, much remains unknown, and more precise estimates of adult infestation prevalence are needed to clarify this relationship.

My objective here is to expand on the findings of Quiroga et al. (2020) by utilizing a species well-suited for investigating *Philornis* parasitism: the Hispaniolan Woodpecker *Melanerpes striatus*. This woodpecker is one of the most abundant birds on Hispaniola, common from sea-level to 2,400 m asl in a wide range of habitats (Latta *et al*. 2006), providing ample sampling opportunities. Additionally, the first *Philornis* species (*P. pici*, reported as *Aricia pici*) was described from a subcutaneous larva collected from an adult Hispaniolan Woodpecker (Macquart 1853). Despite the Hispaniolan Woodpecker’s high abundance, Quiroga et al. (2020) reported only two new records of *Philornis* infestation on adults: one each from the Cordillera Central (prevalence = 20%, *N* = 5 individuals; H.M. Garrod pers. comm.) and Punta Cana (prevalence = 7%, *N* = 14 individuals; L. Soares and S.C. Latta pers. comm.). Furthermore, the parasite negatively impacts the reproductive success of at least one other Hispaniolan endemic, the critically endangered Ridgway’s Hawk *Buteo ridgwayi* (Hayes *et al*. 2019). Yet the woodpecker’s continued abundance in spite of *Philornis* and anthropogenic pressures (Mitchell & Bruggers 1985) suggests it could be an excellent model system to advance *Philornis* biology. To that end, I test three hypotheses (Table 1) regarding *Philornis* infestation prevalence and intensity on adult birds.

**Table 1.**
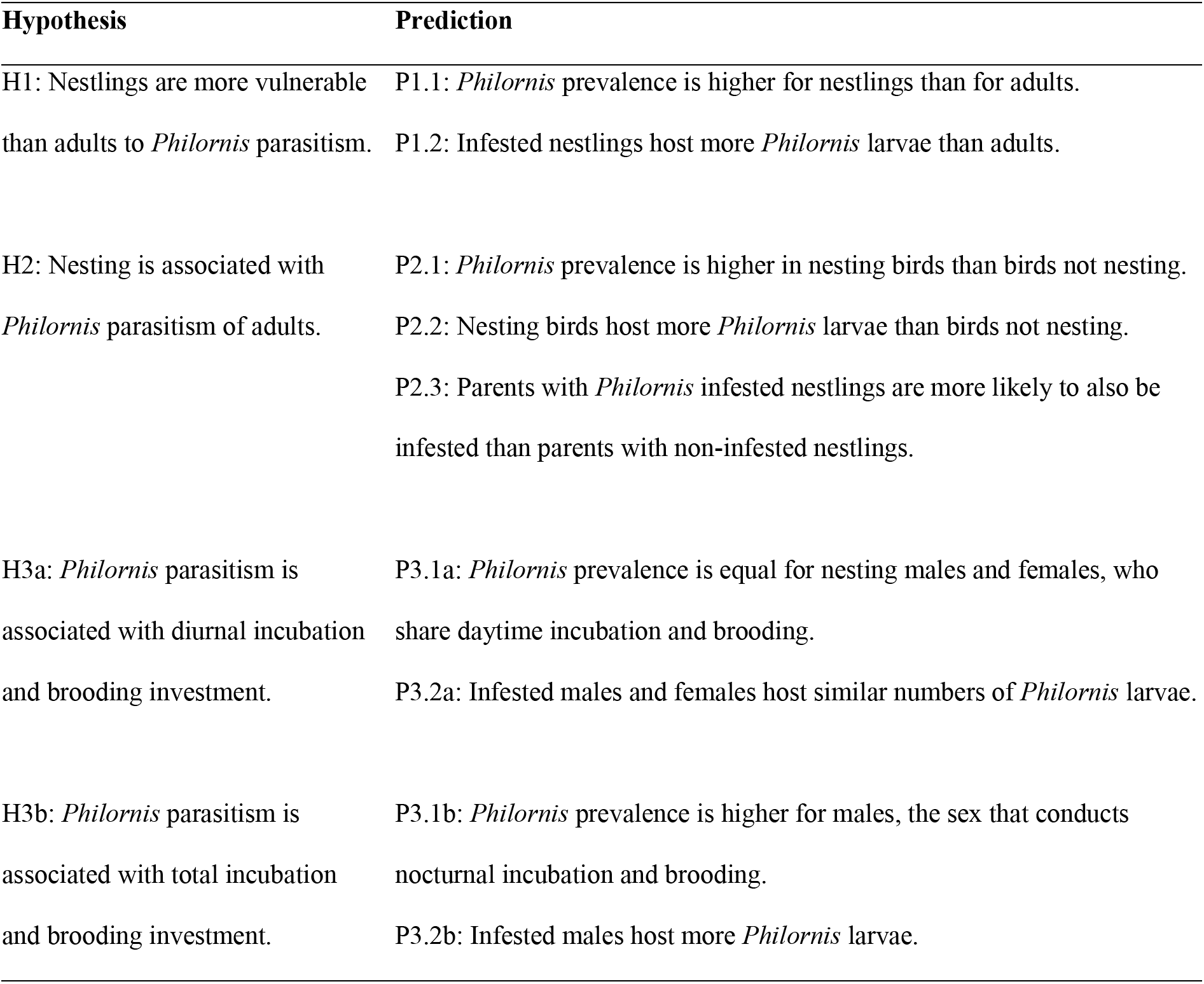
Summary of hypotheses and predictions regarding *Philornis* infestation status.

First, I test two predictions of the hypothesis (H1) that nestlings are more vulnerable than adults to *Philornis* parasitism (Teixeira 1999). This hypothesis predicts that (P1.1) *Philornis* prevalence (the proportion of birds infested) should be higher for nestlings than for adults. Assuming nestlings are easier targets for infestation, this hypothesis also predicts (P1.2) that nestlings should have higher intensity (number of larvae per infested individual) *Philornis* infestations compared with adult birds. Due to their mobility, adult woodpeckers should provide not only fewer opportunities for larval deposition by adult flies across adults, but also fewer opportunities for repeat deposition on individual adults.

Second, I test three predictions of the hypothesis (H2) that nesting behavior itself is associated with *Philornis* parasitism of adults. If *Philornis* is more prevalent and intense on nestlings than adults (Arendt 1985a), parasitism of adults might be an opportunistic direct result of nesting activity. This hypothesis thus predicts that *Philornis* (P2.1) prevalence and (P2.2) intensity should be higher for nesting birds than birds not nesting. This hypothesis also predicts concurrent infestation of parent and nestling birds from the same nest. In other words, (P2.3) parents of infested nestlings should themselves be more likely to be infested than parents of non-infested nestlings.

Lastly, I test four predictions of the hypothesis (H3a,b) that adult *Philornis* parasitism is associated with incubation and brooding investment (Teixeira 1999). While nesting itself might increase exposure to *Philornis*, intersexual differences in breeding behavior might result in females and males experiencing different levels of parasitism. Hispaniolan Woodpeckers are socially and genetically monogamous (LaPergola & Riehl 2022), and both females and males develop brood patches and share approximately equivalent diurnal incubation and brooding (unpubl. data). If incubation and brooding behavior increase exposure (H3a), *Philornis* (P3.1a) prevalence and (P3.2a) intensity should be similar in female and male Hispaniolan Woodpeckers. Like most woodpecker species (Winkler *et al*. 1995), though, male Hispaniolan Woodpeckers perform all overnight incubation of eggs and brooding of nestlings (pers. obs.), a form of essential parental care. This male-biased nocturnal incubation and brooding behavior might be important because adults of at least some *Philornis* species will visit nests at night (O’Connor et al. 2010) and in the late afternoon and dusk (Pike et al. 2021). If overnight incubation and brooding increase exposure (H3b), *Philornis* (P3.1b) prevalence and (P3.2b) intensity should be higher for nesting males than nesting females.

Tests of these hypotheses and predictions (Table 1), which require data from both nestlings and adults from the same population, have only been reported for the Caribbean endemic Pearly-eyed Thrasher *Margarops fuscatus* (Arendt 1985a). Both Pearly-eyed Thrashers and Hispaniolan Woodpeckers nest in cavities, a life-history trait that could impact parasitism exposure (Nilsson 1986), so one might predict similar patterns of *Philornis* prevalence and intensity in both species. In support of H1, nestling Pearly-eyed Thrashers exhibited a far higher prevalence (96%) and intensity (mean = 37 larvae/nestling) of *P. deceptivus* compared with adult prevalence (31%) and intensity (mean = 3.1 larvae/adult) on Puerto Rico (Arendt 1985a). To the best of my knowledge, H2 has not been directly tested in Pearly-eyed Thrashers and has only indirect support from immunological data in the Galápagos endemic Medium Ground Finch *Geospiza fortis*, which showed higher *Philornis*-specific antibody levels during nesting than pre-nesting (Huber *et al*. 2010). Pearly-eyed Thrasher data support H3a since *Philornis* prevalence among nesting females, which perform all incubation and brooding, was ~3.5 times higher than for nesting males (Arendt 1985a). Indirect evidence supporting H3a was also found in the medium ground finch: nesting females, who brood nestlings, had higher *Philornis*-specific antibody levels than nesting males (Huber *et al*. 2010). However, no studies have investigated *Philornis* in a species where males perform essential incubation and brooding.

## METHODS

### Field methods

I studied Hispaniolan Woodpeckers in the community of Piedra Blanca (19.1193°N, 70.5819°W; 550–700 m asl), 3 km east of Jarabacoa, La Vega, Dominican Republic, between April 2012 and July 2017. The site (~84 ha) comprised several private properties on a landscape of pine (*Pinus occidentalis* and *P. caribaea*) and broadleaf wet forest fragments immersed in a matrix of cattle pastures with isolated or clustered royal palms *Roystonea hispaniolana*, small fragments of secondary vegetation, and “living tree” (predominantly *Gliricidia sepium*) fences. This region experiences a mild, dry winter (January–March), followed by a short wet spring season (April–May), a long, dry summer season (June - September), and a short, wet fall season (October–December) coinciding with the latter half of the Atlantic hurricane season (Climate-data.org 2021). Although the Hispaniolan Woodpecker is thought to breed year-round in parts of its range (Latta et al. 2006), the study population exhibits a defined breeding season that lasts six months, spanning March through August with peak clutch initiation in May (unpub. data). This population has nestlings for nearly 160 days of the year, with hatching observed as early as 13 March and as late as 9 August (Fig. S1). For the remainder of the Methods, I use “we” in lieu of “I” to describe most activities because they involved a team of tireless volunteer field assistants.

We evaluated *Philornis* infestation status on nestling and adult woodpeckers at trees monitored for nesting activity, which we selected based on the presence of cavities and nesting activity (e.g., cavity excavation, adults entering/exiting cavities, etc.). To determine nesting activity, we inspected cavities using a penlight and small inspection mirror (1–2” diameter) while climbing or with a wireless camera attached to a 15.2 m telescoping pole that broadcasted images to a portable digital television (Huebner & Hurteau 2007, Waldstein 2012). Once we detected a nesting attempt (i.e., ≥1 eggs), we typically checked the clutch every 3–5 days and, when possible, daily if we did not know the clutch completion date. Incubation typically lasted 11 days (range = 9–14 days). The nestling sampling protocol differed slightly in timing across years, but in general, sampling involved collecting morphometric measurements and inspecting the entire body surface for the presence/absence of *Philornis*, including counting the number of active and empty wounds (Fig. 1). We considered a wound active if it contained ≥1 subcutaneous *Philornis* larvae, and, in cases where ≥2 larvae inhabited the same wound (see Fig. 1a for example of two sets of posterior spiracles of larvae visible in a single wound), we recorded the total number of detectable larvae. Empty *Philornis* wounds resembled active wounds in appearance, except that empty wounds tended to look less swollen (Fig. 1b), lacked detectable larvae, and retained an opening where a larva had resided. For all years when we did not know the nest’s hatch date (e.g., nest was found with nestlings), we sampled and banded nestlings as soon as they were large enough to carry four bands—two colour bands on one leg and one colour band and one metal band on the other leg. For nests with known hatch dates from 2013–2015, we sampled and fully banded nestlings when they were ~14 days old and resampled at ~21 days old. For nests with known hatch dates in 2016 and 2017, we sampled and metal banded nestlings at ~7 days old, resampled at ~14 days old, and resampled and added three colour bands at ~21 days old.

**Figure 1.**
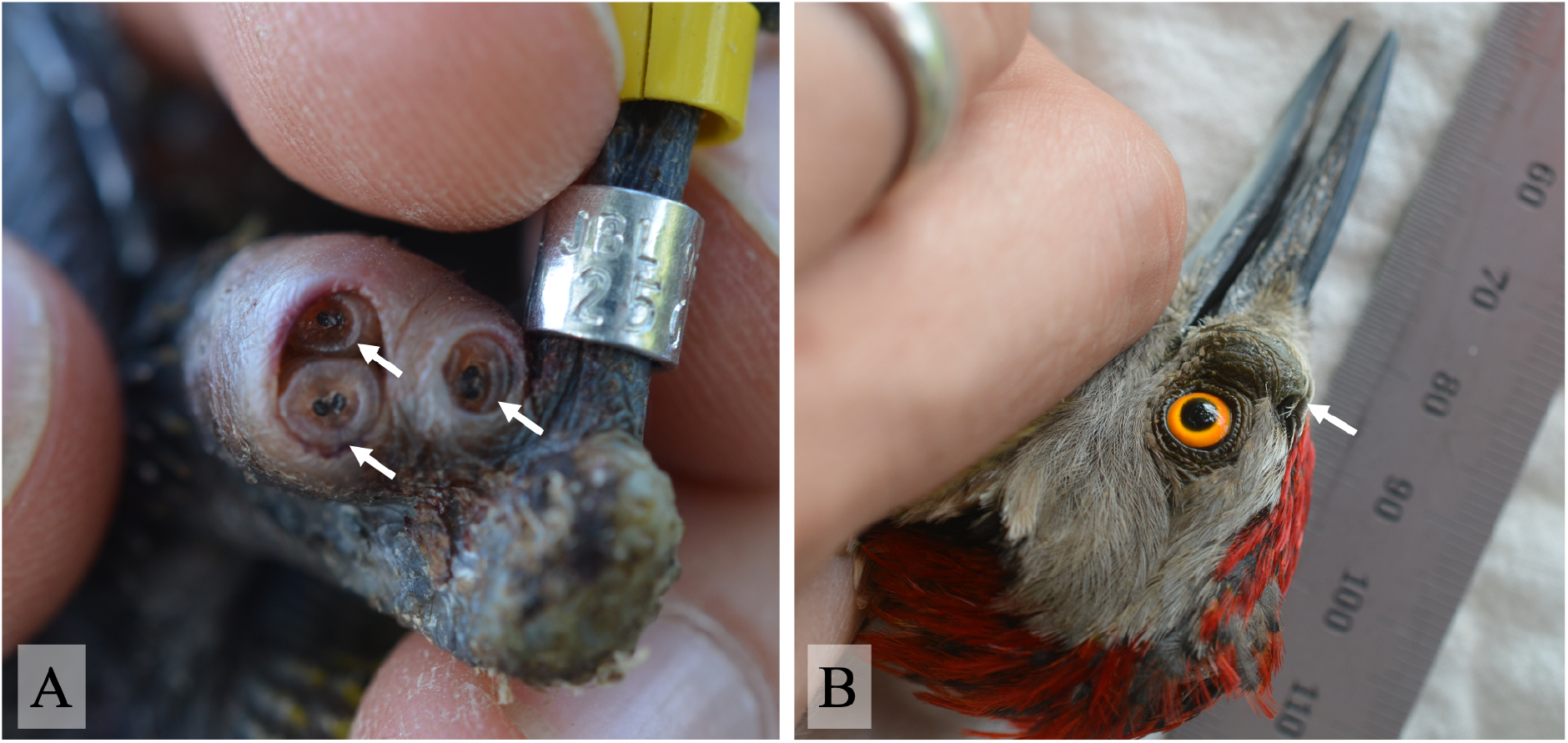
Example of active and empty wounds associated with *Philornis* parasitism in Hispaniolan Woodpeckers. (A) Active wounds containing three *Philornis* larvae (indicated by white arrows) on a nestling woodpecker’s leg, where the posterior spiracles of larvae are visible. Two larvae inhabit one wound while one larva inhabits an adjacent wound. (B) Empty *Philornis* wound on an adult male woodpecker’s face. All photos by author.

For adult sampling, we captured birds via two approaches: (1) ambushing adults in nest cavities and (2) an elevated, dual-tower mist-net system (LaPergola & Kenyon in prep.). Ambushing involved setting up ambush traps as in Stanback and Koenig (1994) to allow pre-dawn capture of roosting birds. See Garrod and LaPergola (2018) for more details on implementation. To reduce nest abandonment, we used the ambush method ≥7 days before egg-laying or ≥22 days post-hatch. The mist-net tower system involved erecting two 15.2 m tower poles supported with guy lines (ropes) and, using pulleys and ropes, raising two stacked 12 m mist-nets in front of nesting trees. This method reduced disturbance at nests, enabled capture of woodpeckers using trees too unstable to climb, increased sampling efficacy before nesting, and increased sampling of non-nesting birds. As with nestlings, each adult received a unique four-band combination and was inspected for the absence or presence of *Philornis*. When present, we counted the number of active and empty *Philornis* wounds. We also recorded sex of adults based on crown colour, which is black for females and red for males.

We determined the nesting and breeder status of captured birds by monitoring nesting attempts and identifying attendant birds via focal nest watches. Nest watches involved 2 or 3 hour sessions in which an observer sat 15–20 m from a nest tree in a burlap blind, trained a 15X or 20X spotting scope on a focal cavity entrance for a nest, and recorded the identities (i.e., band combinations) and behaviors of woodpeckers that visited the nest. For testing the hypothesis that *Philornis* parasitism in adults is associated with nesting, I coded adults as belonging to one of two categories: nesting or not nesting. I counted an adult as nesting if it met two criteria: (1) we observed the bird incubating at or provisioning ≥1 nest within the year of capture, and (2) we captured the bird after the earliest possible clutch initiation date for its earliest possible nesting attempt within the year of capture. I counted banded birds as not nesting if they met one of the following criteria: (1) we captured the bird early in the field season before most nests were initiated (between January and before early April), or (2) the bird was not associated with a nesting attempt prior to the date of capture in the same calendar year.

Although we did not attempt to identify larvae to species, *Philornis pici* is the only *Philornis* species currently known to infest birds on Hispaniola, and, as mentioned earlier, was first described from the Hispaniolan Woodpecker (Macquart 1853). Elsewhere in the Dominican Republic, researchers have confirmed this species to parasitize Ridgway’s Hawk (Hayes *et al*. 2019, Quiroga *et al*. 2020). However, *P. porteri* has also been identified parasitizing Ridgway’s Hawk (M.A. Quiroga pers. comm.). The distribution of *P. porteri* on Hispaniola is currently unknown, but it is possible that the *Philornis* detected in the present study could be *P. pici*, *P. porteri*, or both.

### Statistical analyses

See Table 1 for a summary of the hypotheses and their predictions. For testing the hypothesis that nestlings are more vulnerable to *Philornis* parasitism than adults (H1), I tested the two predictions with separate generalized linear mixed-effects models (GLMMs). For the prediction that the probability of being parasitized is higher for nestlings than for adult birds (P1.1), I used a GLMM with a binomial fit to test for an association between infestation status and age coded as a categorical fixed effect (adult vs. nestling). Infestation status was treated as a binary response (0 = no evidence of *Philornis*, 1 = presence of ≥1 *Philornis* larvae, empty wounds, or both) in this model. For the prediction that nestlings host greater numbers of *Philornis* wounds than adults (P1.2), I used a GLMM with a negative binomial distribution to test for an association between the total number of *Philornis* wounds (summing the numbers of empty and active wounds, or total number of larvae) and age. Because many birds were never observed with infestations, including all sampled individuals would lead to zero-inflation for the total number of *Philornis* wounds; consequently, I used a manual hurdle model approach, including only infested birds in this model.

To test the predictions of the hypotheses that (H2) nesting and (H3a, H3b) incubation and brooding investment are associated with *Philornis* parasitism in adults, I used four GLMMs to test for associations of adult infestation status with nesting status, sex, and the interaction of nesting status and sex. I coded both nesting status (nesting vs. not nesting) and sex (female vs. male) as categorical fixed effects for all four models. To test predictions regarding prevalence (P2.1, P3.1a, and P3.2a), I used GLMMs with a binomial fit: infestation status was treated as a binary response as with the analysis comparing nestlings and adults. For the first GLMM, I included all adults of known nesting status, retaining birds known not to have bred within the year of capture. For the second GLMM, I retained only birds known to have bred within the year of capture to exclude any effects of unanticipated differences between breeders and non-breeders. This more restrictive analysis contrasted known breeders captured prior to their first nesting attempt within the breeding season (“not nesting”) with breeders actively nesting at the time of capture (“nesting”). For testing the predictions regarding intensity as they relate to nesting status and sex (P2.2, P3.2a, and P3.2b), I used a GLMM with a Poisson distribution to test for an association of the total number of *Philornis* wounds (summing the numbers of active and empty wounds, or total number of larvae) with nesting status, sex, and the interaction effect of nesting status and sex. As with the nestling-adult comparison, many observations involved no infestation and would lead to zero-inflation for the total number of *Philornis* wounds; consequently, I used a manual hurdle model approach, including only infested adults in this model. Additionally, I only ran this model with the dataset that included all adults of known nesting status, including birds known not to have bred within the year of capture.

To test the prediction (P2.3) that parents with *Philornis* infested nestlings are themselves more likely to also be infested than parents with non-infested nestlings, I analyzed the subset of sampled parent birds whose nestlings were also sampled. I used two Fisher’s exact tests because sample sizes were insufficient to accommodate a GLMM approach, and I restricted analyses to the level of the nest to avoid pseudoreplication. First, I compared the proportion of nests with at least one infested adult based on the presence of any *Philornis* wounds for nests in which one or more nestlings had any *Philornis* wounds (i.e., infested) with nests in which nestlings remained free of *Philornis* (i.e., non-infested). Using this same set of nests, I made a second comparison of the proportion of nests with at least one adult bearing only active *Philornis* wounds.

For all models, I also included capture date as a continuous fixed effect based on the following. The capture date range was fairly large (range = 168 d, 28-Feb–4-Aug), which included the end of the winter dry season, the short wet spring season, and the long dry summer season. Furthermore, previous studies have documented a positive association between the probability of adults and nests having *Philornis* and the timing of breeding (Arendt 1985a, Rabuffetti & Reboreda 2007). For all analyses, I scaled capture date in day of year format via Z-transformation by subtracting the mean capture date and dividing by the standard deviation.

For all models except for those testing predictions regarding intensity only in adults (P2.2, P3.2a, and P3.2b), I included the following as random effects: the tree where a bird was captured or, for known breeders, where it bred in the year of capture (Tree ID); year of capture; and individual ID. I included Tree ID as a random effect because the Hispaniolan Woodpecker is unique among the Picidae, being one of only three known woodpecker species to exhibit facultative colonial nesting. Within the same population, Hispaniolan Woodpeckers pairs can nest singly or in clusters, with two or more pairs nesting concurrently in separate cavities on the same tree (Short 1974, Winkler *et al*. 1995, LaPergola 2018). Additionally, I wanted to account for the non-independence of nestlings from the same brood and thus the same parents, but using a nest ID random effect would have precluded using adults without nests. Using Tree ID as a random effect is thus a more conservative approach to account for non-independence, especially for nestlings. I included year as a random effect in all analyses because I was not confident that interannual variation was sampled adequately enough to interpret the fixed effects of year (Bennington & Thayne 1994). Lastly, I included individual ID because some individuals were captured multiple times. For testing predictions regarding intensity only in adults (P2.2, P3.2a, and P3.2b), I included only year as a random effect because including Tree ID and individual ID led to failed model convergence.

I conducted all statistical analyses in RStudio v. 1.1.463 using R v. 3.6.3 (R Core Team 2020). For fitting GLMMs, I used the *glmer* (binomial and Poisson fits) and *glmer.nb* (negative binomial) functions in the *lme4* package (Bates *et al*. 2015). I used the *fisher.test* function for Fisher’s exact tests. For models where interaction terms were not significant, I report only the results of the additive models. All means are reported ± the standard error of the mean, and all confidence intervals for count data are 95% and were calculated via the Wald Method.

## RESULTS

### Summary of *Philornis* parasitism prevalence

Over six years, I obtained 218 adult records representing 184 unique individuals (83 females and 101 males), which included 26 individuals (eight females and 18 males) recaptured once and four individuals (one female and three males) recaptured twice. Of all adult records, 40 (18%; CI = 14–24%) included individuals with evidence of *Philornis* parasitism. Of all individuals (*n* = 184), 36 (20%; CI = 14–26%) had evidence of *Philornis* parasitism, which included 24 (67%; CI = 50–80%; *n* = 36 individuals) with *Philornis* empty wounds, nine (25%; CI = 14–41%) with active wounds, and three (8%; CI = 2–23%) with both empty and active wounds. Of all adults with more than one capture (*n* = 26), 11 individuals exhibited changed infestation status (Table S1). These records included four individuals recaptured within the same year, of which two had active wounds on the second capture but no wounds on the first, one had old wounds on the first capture but not the second, and one individual had old wounds on the first capture but no visible wounds on the second capture 82 days later.

Across six years, I collected 554 nestling records representing 381 individuals from 127 nesting attempts. These figures amounted to a mean of 4.4 ± 2.4 records per nesting attempt (range = 1–10 records per nesting attempt) and a mean of 3.0 ± 1.0 nestlings per nesting attempt (range = 1–5 nestlings per nesting attempt). Of all nestling records, 123 (22%; CI = 19–26%) showed evidence of *Philornis* parasitism, and of all nestlings observed, 107 (28%; CI = 24–33%) exhibited evidence of *Philornis* parasitism on at least one sampling event. Of the nestling individuals with evidence of *Philornis*, most (73%; CI = 64–80%) involved active wounds (45 observations with only active wounds and 33 observations with both active and old wounds), while fewer observations involved only old wounds (19%; CI = 12–27%; for 8% of nestling observations, the wound status was not recorded). Infested nestlings came from 43 (34%; CI = 26–43%) of all monitored nesting attempts.

### H1: Comparison of adults and nestlings

Using the full set of adult and nestling capture records, age and scaled day of year captured alone were not significant predictors of the presence/absence of *Philornis* parasitism (Table 2a). However, there was a significant interaction effect of age and scaled day of year captured, such that the probability of exhibiting *Philornis* parasitism increased with the scaled day of year for nestlings but not for adults (Fig. 2A). In contrast to presence/absence, age alone was significantly associated with the total number of *Philornis* wounds (empty plus active wounds) (Table 2b). Infested nestlings had an average of 7.1 ± 0.5 *Philornis* wounds (range = 1–39 *Philornis* wounds; *n* = 123 nestling records) while infested adults had an average of only 2.0 ± 0.2 wounds (range = 1–5 *Philornis* wounds; *n* = 40 adult records; Fig. 2b).

**Figure 2.**
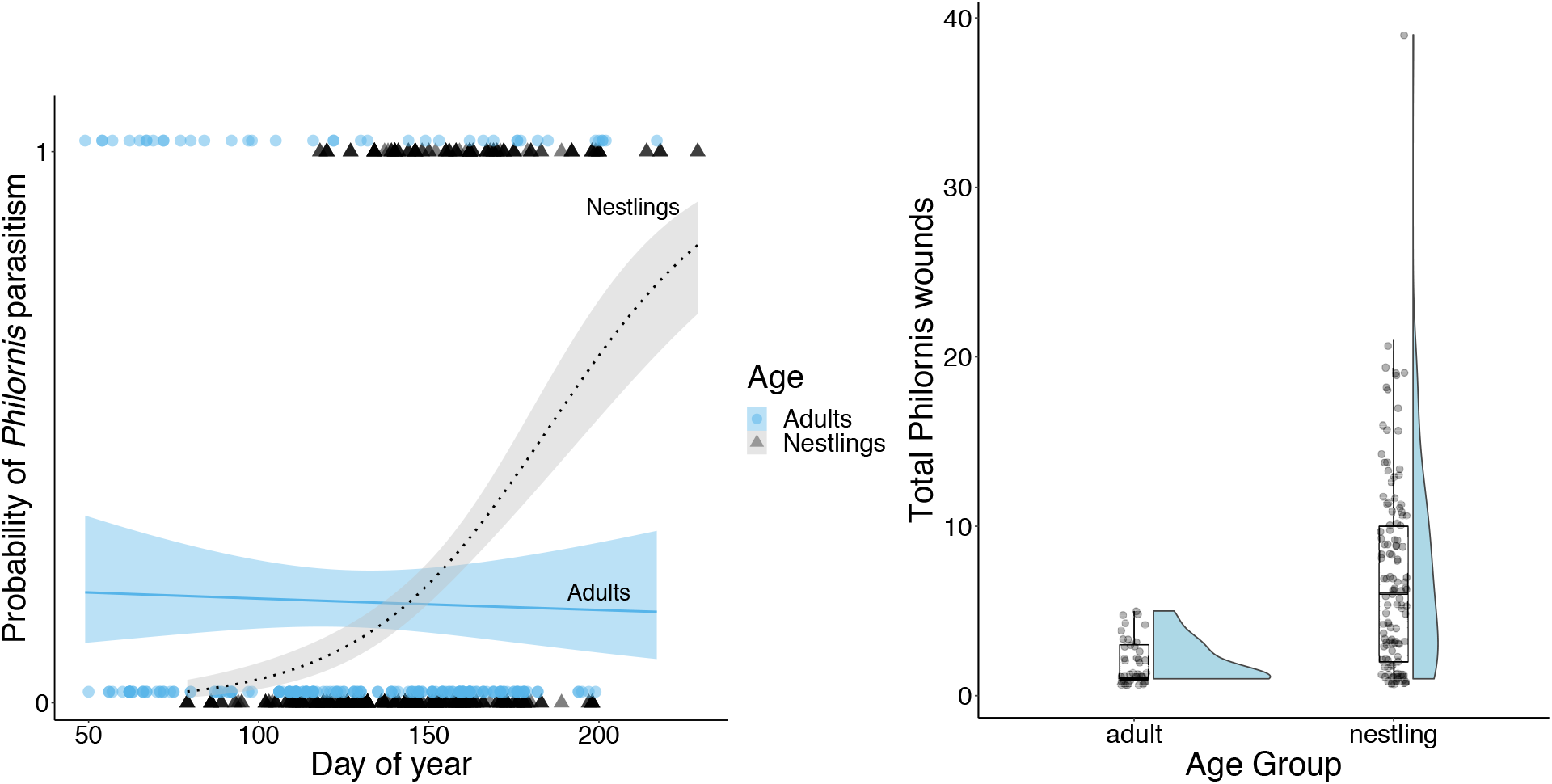
Probability and intensity of *Philornis* parasitism on adult and nestling Hispaniolan Woodpeckers. (A) Probability of *Philornis* parasitism plotted as raw data (adults = blue circles, *n* = 218 observations; nestlings = gray triangles, *n* = 554 observations) and model predictions from a generalized linear mixed model testing for an association with age, day of year captured, and their interaction. The blue solid line and black dashed line represent model predictions for adults and nestlings, respectively. Raw data were artificially vertically separated to improve visibility of points. (B) Raincloud plot comparing adults (*n* = 40 observations) and nestlings (*n* = 123 observations) for the total number of *Philornis* wounds observed on infested individuals (i.e., only non-zero values for the total number of *Philornis* wounds). Sample sizes indicate the number of observations.

**Table 2.** Results of two generalized linear mixed-effects models testing for an association of *Philornis* parasitism with age in Hispaniolan Woodpeckers. Model (a) included the binary response of *Philornis* parasitism (yes/no) and fixed effects of age (adult vs. nestling), scaled date of capture (DOY scaled), and their interaction, and was fit with a binomial distribution. Model (b) tested for an association of total number of *Philornis* wounds on infested birds only with age, DOY scaled, and their interaction, and was fit with a negative binomial distribution. Random effects for both models were individual identity (a: *n* = 559 individuals; b: *n* = 143 individuals), year of capture (*n* = 6 years in both a and b), and tree ID where captured or bred (a: *n* = 41 trees; b: *n* = 25 trees).

| Model and factors | Estimate ± S.E. | z-value | P |
| --- | --- | --- | --- |
| a. *Philornis* parasitism (yes/no) |  |  |  |
| Intercept | -3.092 ± 0.971 | -3.184 | 0.00145 ** |
| Age (nestling) | -0.689 ± 0.365 | -1.888 | 0.059 |
| DOY Scaled | 0.170 ± 0.205 | 0.829 | 0.407 |
| ***Age (nestling) \* DOY Scaled*** | **2.383 ± 0.438** | **5.443** | **5.25e-08 \*\*\*** |
| b. Total *Philornis* wounds |  |  |  |
| Intercept | 0.598 ± 0.192 | 3.113 | 0.00185 ** |
| ***Age (nestling)*** | **0.935 ± 0.191** | **4.895** | **9.82e-07 \*\*\*** |
| DOY scaled | -0.004 ± 0.085 | -0.047 | 0.962 |

### H2 and H3: Nesting status and sex

When restricting the analyses to adults of known nesting status, there was no significant association between *Philornis* infestation and whether an adult was currently nesting (Table 3). This result was true for both the analysis including all adults of known nesting status, i.e., retaining birds known not to have bred within the capture year (Table 3a), and for the analysis restricted to only birds that nested within the capture year (Table 3b). Additionally, the scaled day of year was not significantly associated with infestation status. There was no significant interaction between sex and nesting status, but adult sex was significantly associated with infestation in both analyses. For all adults of known nesting status, 9% of females (CI = 4–17%; *n* = 82 observations) and 27% of males (CI = 20–36%; *n* = 111 observations) showed signs of current or past infestation (Figure 3). These proportions remained similar for the subset that included only birds that nested within the capture year (8% of females: CI = 4–18%, *n* = 60 observations; 26% of males: CI = 17–37%, *n* = 74 observations).

**Figure 3.**
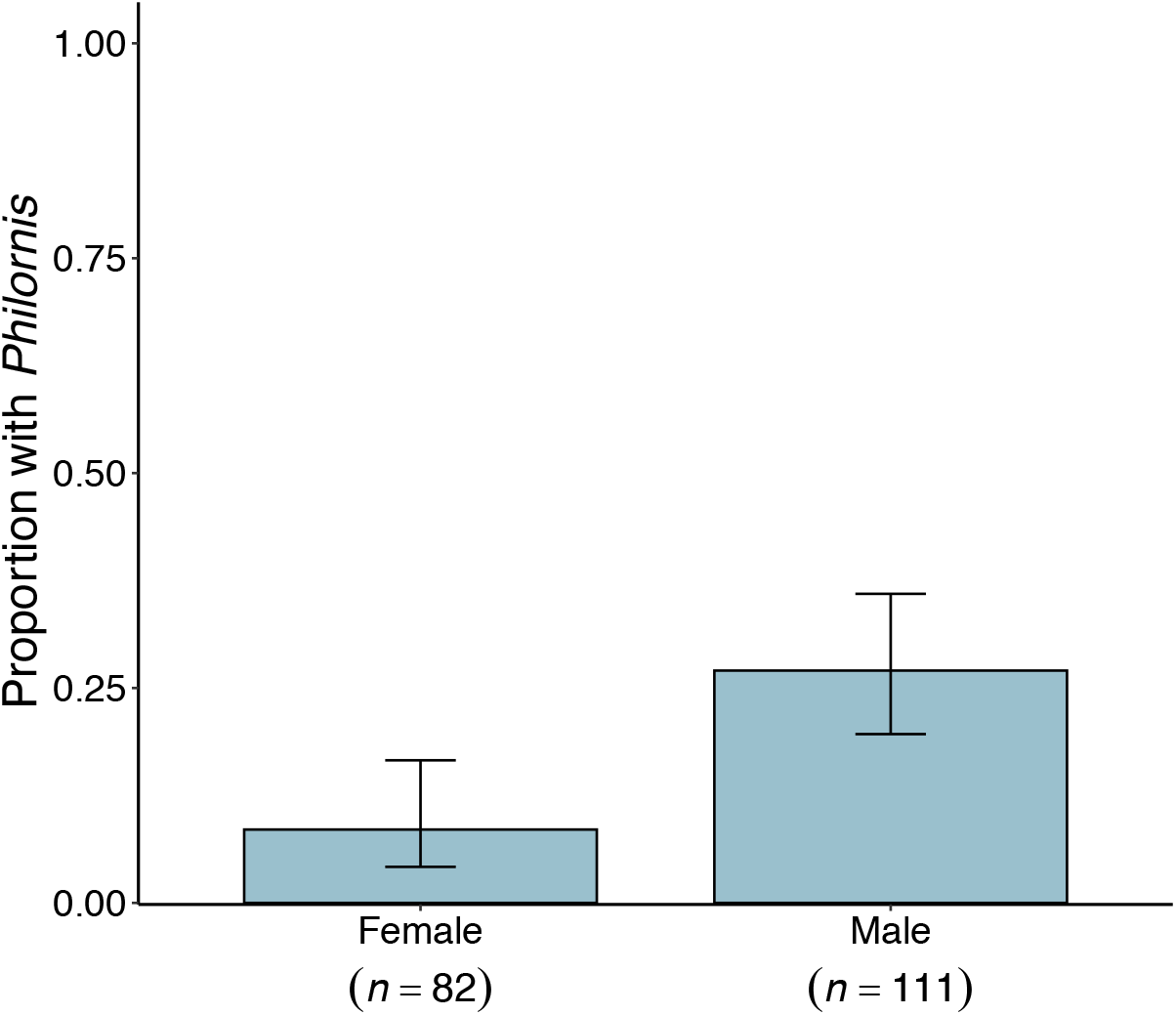
Proportion of female and male adult Hispaniolan Woodpeckers with ≥1 *Philornis* parasite. Male woodpeckers were significantly more likely to be infested (Table 3). Error bars represent 95% confidence intervals. Sample sizes indicate the number of observations.

**Table 3.** Results of two generalized linear mixed-effects models testing for an association of the binary response of *Philornis* parasitism (yes/no) of adult Hispaniolan Woodpeckers with sex (female or male), nesting status (actively nesting or not nesting at time of capture), and the scaled day of the year captured (DOY scaled). Random effects were individual identity (a: *n* = 163 individuals; b: *n* = 113 individuals), tree ID where an individual was captured or bred (a: *n* = 30 trees; b: *n* = 27 trees), and year of capture (*n* = 6 years in both a and b). Model (a) included the full set of individuals with known nesting status within a year, including individuals that never bred (non-breeders) within the capture year (*n* = 193 observations). Model (b) included only individuals that were known to have bred within the capture year (*n* = 134 observations).

| Model and factors | Estimate ± S.E. | z-value | P |
| --- | --- | --- | --- |
| a. *Philornis* parasitism (yes/no) on breeders and non-breeders |  |  |  |
| Intercept | -3.116 ± 0.874 | -3.565 | 0.0004 |
| ***Sex (male)*** | **1.674 ± 0.483** | **3.468** | **0.0005\*\*\*** |
| Nesting Status (not nesting) | 0.312 ± 0.651 | 0.480 | 0.632 |
| DOY Scaled | 0.091 ± 0.307 | 0.297 | 0.767 |
| b. *Philornis* parasitism (yes/no) on breeders only |  |  |  |
| Intercept | - 3.391 ± 0.961 | -3.529 | 0.0004 |
| ***Sex (male)*** | **1.697 ± 0.592** | **2.869** | **0.0041\*\*** |
| Nesting Status (not nesting) | 1.221 ± 0.898 | 1.360 | 0.174 |
| DOY Scaled | 0.475 ± 0.384 | 1.238 | 0.216 |

Infested female and male adults had similar numbers of *Philornis* wounds. Infested females had a mean of 2.0 ± 0.7 wounds (range = 1–5; *n* = 7 observations), and infested males had a mean of 2.0 ± 0.2 (range = 1–5; *n* = 29 observations). None of the fixed effects were significant in the model (Table 4).

**Table 4.** Results of a generalized linear mixed-effect model with a Poisson distribution testing for an association between the total number of *Philornis* wounds on adult Hispaniolan Woodpeckers and sex (female or male), nesting status (actively nesting or not nesting at time of capture), the interaction of sex and nesting status, and scaled day of year captured (DOY scaled), including year as a random effect (*n* = 4 years). This analysis used birds of known nesting status (*n* = 36 observations), including birds that bred and those that did not within the year of capture.

| Factors | Estimate ± S.E. | z-value | P |
| --- | --- | --- | --- |
| Intercept | 0.601 ± 0.357 | 1.684 | 0.0921 |
| Sex (male) | 0.045 ± 0.230 | 0.150 | 0.8804 |
| Nesting Status (not nesting) | 0.100 ± 0.454 | 0.220 | 0.8256 |
| DOY Scaled | -0.126 ± 0.232 | -0.543 | 0.5874 |

The 12 adults with active infestations were mostly (58%) known breeders (Table S2). Two of the three females with active infestations also had infested nestlings at the time of capture. Of the nine males with active infestations, four had infested nestlings at the time of capture and one male had fledged two young (one infested and one not) one month prior to capture. The remaining individuals of uncertain breeding status (one female and four males) were all caught within the known breeding season at the site (Fig. S1); the earliest capture was on 8 March and the latest capture 20 July.

Analyses of only adults for which infestation status of nestlings was known (*n* = 41 nests representing 40 unique parents or parent pairs) provided evidence of concurrent infestation of adults and young. Of nests with infested young, 53% (95% CI = 30–74%, *n* = 17) also had at least one parent infested while only 17% (95% CI = 6–36%, *n* = 24) of nests with non-infested young had at least one parent infested (Fisher’s exact test: <u>*P* = 0.017</u>). When considering only active *Philornis* wounds on adults, 29% (95% CI = 13–53%, *n* = 17) of nests with infested young also had at least one infested parent while 0% (95% CI = 0–16%, *n* = 24) of nests with non-infested young had infested parents (Fisher’s exact test: *P* = 0.008).

## DISCUSSION

Most previous work on *Philornis* myiasis has understandably focused on these parasites’ impacts on nestling birds (Arendt 1985b, Dudaniec & Kleindorfer 2006, Hayes *et al*. 2019) since this life stage has long been considered the primary target of parasitism (Teixeira 1999). The present study is one of very few that has concurrently documented *Philornis* prevalence and intensity on nestlings and adults from the same population (see also Arendt 1985a). Intriguingly, adult Hispaniolan Woodpeckers were just as likely to exhibit evidence of *Philornis* as nestlings, although nestlings experienced greater intensity of infestation. Among adults, while nesting status itself was not a significant predictor of being infested, nests with infested nestlings were significantly more likely to also have one or both parents be infested than nests with non-infested nestlings, and males, which invest more in overall incubation and brooding, were significantly more likely to have *Philornis* infestations than females.

### Nestlings vs. adults

The present study’s results falsify the first prediction of the hypothesis that nestlings are more vulnerable to *Philornis* parasitism but support the second prediction that nestlings experience more intense infestations. While adult and nestling Hispaniolan Woodpeckers did not differ in the probability of being parasitized (Table 2a), the probability of being parasitized for nestlings did increase with passage of the breeding season yet remained more or less static for adults across the breeding season (Fig. 2a). Furthermore, when infested, nestlings bore greater numbers of *Philornis* wounds than did adult birds (Table 2b; Fig. 2b.). This difference in intensity is likely due to the increased accessibility of nestlings to *Philornis* flies in contrast to lower accessibility of adults. Of the two results and corresponding predictions, the contrast in *Philornis* intensity supports the hypothesis that nestlings are indeed more vulnerable to parasitism. However, the similarity in prevalence suggests a complementary hypothesis that adult *Philornis* are equally likely to find nestling and adult Hispaniolan Woodpeckers but that nestlings are less resistant to infestation. Unfortunately, we currently lack the necessary *Philornis* natural history data to evaluate this possibility. If *Philornis* females oviposit directly in woodpecker nests, this behavior would help explain higher intensity of *Philornis* on nestling woodpeckers since adults would have even lower overall exposure to infestation. *Philornis downsi* oviposits in the nest material (Lahuatte et al. 2016), and at least one subcutaneous species, *P. torquans*, will oviposit on inanimate surfaces in captivity (Patitucci et al. 2017, Saravia-Pietropaolo *et al*. 2018). It is therefore plausible, though yet to be confirmed, that *P. pici* and *P. porteri*, the two species most likely parasitizing Hispaniolan Woodpeckers, oviposit directly in the nest.

Regardless of the manner of egg/larval deposition, there are at least three non-mutually exclusive mechanistic hypotheses for lower parasite intensity on adults. First, the adults’ well-developed plumage might reduce accessibility by presenting a physical barrier to burrowing larvae (Oniki 1983). This might be especially relevant for Hispaniolan Woodpeckers as they hatch naked and remain so until 7–8 d post-hatch when their pin feathers typically begin erupting, and while pin break begins around 14 d post-hatch, these feathers fail to cover most of the body other than the feather tracts until about 21 d post-hatch (unpubl. data). Thus the young have little to no physical barrier against *Philornis* other than a brooding adult for roughly their first three weeks. Hispaniolan Woodpecker nestlings also remain in the nest 29–38 d post-hatch (unpubl. data), providing additional exposure time, albeit with an increasing amount of feather coverage. A non-mutually exclusive alternative hypothesis is that the greater mobility of adults reduces their accessibility to *Philornis* (Teixeira 1999). When not actively attending a nest, adults are literally moving targets for flies, covering areas of 1.7–4.2 km^2^ while foraging (Mitchell & Bruggers 1985), whereas nestlings remain relatively stationary, confined to the same nest cavity until fledging. A third hypothesis is that the immune memory of adult birds might make them better able to resist infestation. This immune defense hypothesis is plausible given that mother, but not nestling, Medium Ground Finches had elevated levels of *P. downsi*-binding antibody when exposed to the parasite (Koop et al. 2013). Whether any of these mechanisms can explain differences in *Philornis* infestation intensity for Hispaniolan Woodpeckers remains to be examined.

The patterns of *Philornis* prevalence and intensity reported here for Hispaniolan Woodpeckers contrast somewhat with those from the only other study (see Arendt 1985a) comparing nestlings and adults in the same population. Whereas *Philornis* prevalence among nestling Pearly-eyed Thrashers was much higher than for adults (96% vs. 31%, respectively; Arendt 1985a), prevalence was only non-significantly higher for nestling woodpeckers than for adults (28% vs. 20%). This contrast might be explained by two inter-related factors that differ between the Pearly-eyed Thrasher study and the present study: habitat and climate. The thrasher study took place in a tropical rainforest, with annual rainfall averaging 4460 mm (Arendt 1985a), while the present study occurred in more open, drier habitat, with annual rainfall of 1723 mm (Climate-data.org 2021). Rainfall is a significant predictor of *Philornis* infestation, showing a positive correlation with intensity (Antoniazzi *et al*. 2011, Manzoli *et al*. 2013), and moisture and humidity predict the geographic distribution of at least one *Philornis* species (Cuervo *et al*. 2021). The greater canopy cover and humidity of the rainforest might have promoted larger populations of adult *Philornis* than those in the drier habitat of the Dominican Republic, and these hypothetical larger fly populations might have more fully exploited the vulnerable nestling thrashers while adults could effectively avoid or prevent parasitism. Alternatively, the drier habitat on the Dominican Republic might have reduced access for adult *Philornis* because they would have needed to cross open (i.e., no canopy cover) habitat to reach woodpecker nests. In other words, the Hispaniolan Woodpecker’s habitat structure provides a barrier for adult *Philornis* so they are prevented from fully exploiting the vulnerable nestling woodpeckers. The pattern of intensity differences was similar, though: both nestling thrashers and woodpeckers had greater intensity of *Philornis* than adult birds, and this aligns with the second prediction of the hypothesis that nestlings in both species are more vulnerable to *Philornis* parasitism.

One limitation of the present study was that the precise timing of active infestation for adults was often unknown, especially relative to timing of nesting. This issue arose because most evidence of *Philornis* on adult Hispaniolan Woodpeckers was in the form of empty wounds rather than active wounds containing larvae (33% of adult records involved wounds containing ≥ 1 larvae). In contrast, most nestling observations involved active wounds. This difference is due in part to the sampling effort relative to the age of target birds. Nestling Hispaniolan Woodpeckers were sampled at known ages and within 25 days of hatching so the period of exposure was limited, increasing the probability of detecting subcutaneous *Philornis* larvae, which can remain attached for 5–8 days (Arendt 1985a, Young 1993). The exposure period for adult birds bearing empty wounds, however, was presumably all the days they lived prior to the date of capture, decreasing the probability that we would detect their wounds when they contained larvae. This limitation is important for two reasons regarding timing. First, while it might be most parsimonious to assume all adult Hispaniolan Woodpeckers with empty *Philornis* wounds were infested as adults, we do not know the maximum number of days empty wounds persist after larval detachment from woodpeckers. This uncertainty means that some adults bearing empty wounds might have been infested as nestlings though this seems unlikely. Quiroga et al. (2020, p. 2) posited that all adults in their sample were likely parasitized as adults because “scars [i.e., empty wounds] usually heal ca. one week after larvae detach from the host…”. There are few published accounts of the time it takes for an empty wound from a subcutaneous *Philornis* infestation to heal completely and leave no visible trace, but scars left by subcutaneous *Philornis* after removal from nestling hosts of three species (Baywings *Agelaioides badius;* Screaming Cowbirds *Molothrus rufoaxillaris;* and Shiny Cowbirds *M. bonariensis*) lasted at least two days (Ursino et al. 2019). Second, uncertainty of adult exposure potentially reduces the accuracy of nestling-adult seasonality comparisons.

The difference between adults and nestling Hispaniolan Woodpeckers with respect to the seasonality of prevalence begs further consideration. The lack of an effect of day of capture on prevalence in adults might be related to the abovementioned limitation: i.e., sampling date relative to the day(s) of active infestation. Because the majority of nestling observations involved active wounds while most adult records only involved empty wounds, the day of capture for adults was a less reliable indicator of the timing of infestation for them. In other words, it could be that adults showed the same type of seasonality in infestation as nestlings, with the probability of being infested increasing as the season progressed, but the sampling effort precluded detecting such a pattern. If the difference in seasonality between adults and nestlings was a real pattern, though, Hispaniolan Woodpeckers would differ from Pearly-eyed Thrashers, in which adults showed increasing prevalence of *Philornis* as the season progressed but prevalence among nestlings was high throughout the nesting season (Arendt 1985a). To more fully understand the seasonality of *Philornis* infestation will require data on the seasonality of emergence and population dynamics of adult flies (e.g., Causton et al. 2019). Unfortunately, there are no published data on the seasonality of *Philornis* emergence for Hispaniola.

### Nesting status, sex, and brooding/incubation investment

The nesting status of Hispaniolan Woodpecker adults was not significantly associated with prevalence nor intensity of *Philornis* parasitism (Tables 3 and 4), refuting the first two predictions (see P2.1 and P2.2, Table 1) of the hypothesis that such parasitism is associated with nesting. Yet the third prediction (P2.3) of this hypothesis was supported: parents with *Philornis* infested nestlings were more likely to also be infested than parents with non-infested nestlings. Strikingly, only parents of infested nestlings had active wounds whereas none of the parents of non-infested nestlings were observed with active wounds. To the best of my knowledge, these results represent the first direct test of this hypothesis. The lack of an effect of nesting status in the present study could be an artifact of the sampling period, which mostly comprised the nesting season. However, the inclusion of adults known to not be actively nesting at the time of capture should lessen the impact of such an artifact. Another possible limitation was the uncertainty around the time when an adult was first infested because it makes it harder to discern the amount of overlap between infestation and nesting. It will be crucial to more precisely define the window of infestation for sampled adults to accurately compare prevalence and intensity among nesting and non-esting birds in future studies. One could achieve increased accuracy here by sampling more birds in the non-breeding season and capturing more adults when they have chicks of known age. Regarding the latter suggestion, my current sampling, albeit somewhat modest in size (*n* = 41 nests, Table S3), supports the possibility that at least some adults were exposed to infestation when their nestlings were infested. However, it is worth considering whether the observed pattern is not an artifact, i.e., nesting and non-nesting Hispaniolan Woodpeckers are equally likely to be parasitized. If *Philornis* typically finds hosts by searching for nest-related cues (e.g., olfactory), adult Hispaniolan Woodpeckers might be parasitized outside the context of actively breeding if they spend time in nest cavities for other activities. For example, Hispaniolan Woodpeckers roost in previously used nest cavities (pers. obs.). If the cues adult *Philornis* use to find nestlings remain detectable, opportunistic parasitism of adult woodpeckers could occur. Such a scenario might apply in the non-breeding season or even within the breeding season prior to active nesting. One could test this idea experimentally by setting un-baited traps for adult *Philornis* in old or recently used cavities. Another possible reason that nesting status might be less relevant for Hispaniolan Woodpeckers concerns their habit of colonial nesting. For example, adults lacking active nests might still be subjected to parasitism when one or more other colony members are nesting and thus attracting adult *Philornis*. This hypothesis and the impacts of colonial nesting on *Philornis* parasitism more broadly warrant further study since group-living can either increase (Brown & Brown 1986) or decrease (Mooring & Hart 1992) the risk of parasitism. Local heterospecific nesting density was associated with increased intensity of the invasive *P. downsi* (Kleindorfer & Dudaniec 2009), indicating that this hypothesis is well worth investigating in the native ranges of *Philornis* (e.g., see Antoniazzi et al. 2011).

The combined results of adult Hispaniolan Woodpeckers being parasitized regardless of nesting status and nestlings and adults exhibiting similar prevalence suggest that parasitism of adult woodpeckers might be part of a mixed strategy by *Philornis* in which they target adult birds. As suggested by Quiroga et al. (2020), such a strategy might allow flies to reproduce when nestlings are unavailable or in short supply. In the present study, the Hispaniolan Woodpecker population had a defined breeding season, beginning in early March, peaking in May, and tapering off in August (LaPergola 2018, see also Fig. S1) so nestling woodpeckers are unavailable for approximately half the year and only abundant for roughly three months. Some other local species that might host *Philornis* (e.g., *Crotophaga ani, Columbina passerina, Zenaida aurita, Z. asiatica*, and *Coereba flaveola*) have been suggested to breed year-round (Latta *et al*. 2006), but the extent to which they do so in addition to whether they are parasitized at the study site remains unknown. Additionally, capture records of Hispaniolan Woodpeckers from the site are unavailable from most of the non-breeding season, especially September through December. Fully testing this hypothesis that *Philornis* target adult birds when nestlings are unavailable or scarce requires year-round monitoring for infestation of both adults and nestlings and data on the availability and abundance of host nestlings.

Despite the non-significant effect of nesting status, there was some support for the hypothesis that *Philornis* parasitism is associated with incubation and brooding investment in Hispaniolan Woodpeckers. Although the sexes did not differ in intensity of infestation, males were 3.4 times more likely than females to host *Philornis*. This result mirrors the pattern observed in Pearly-eyed Thrashers, where females, who are the sole incubators/brooders, were 3.5 times more likely to host *Philornis* than males (Arendt 1985a). Since female Hispaniolan Woodpeckers perform only diurnal incubation/brooding while males perform diurnal *and* nocturnal incubation/brooding, males might experience increased *Philornis* exposure at night. Unfortunately, almost nothing is known about the temporal activity patterns of *Philornis* on Hispaniola nor for most other *Philornis* with subcutaneous larvae. In the Galápagos, adult *P. downsi* enter host nests to oviposit when the parent birds are absent during the day when nestlings are young and at night when nestlings are older (O’Connor *et al*. 2010), and peak nest visitation rates of adult flies occurs in the late afternoon and dusk in the nestling phase (Pike et al. 2021). But *P. downsi* larvae are free-living and hematophagous and eggs are oviposited in the nest. An important assumption of the hypothesis that nocturnal incubation increases exposure thus needs testing. Additionally, the lack of an interaction effect of nesting status and sex suggests alternative hypotheses warrant testing.

Three major sets of alternative explanations for higher prevalence of *Philornis* among adult male Hispaniolan Woodpeckers are sexual dimorphisms in behavior, morphology, and immunology (Zuk & McKean 1996). One behavioral difference could be that males experience greater exposure by spending more time in a particular site or habitat (e.g., Tinsley 1989). For example, male Hispaniolan Woodpeckers might spend more time than females in cavities overall, even when not tending a nest with eggs or young. This could be the case if males played a larger role in defending cavities from competitors and were thus more likely to encounter *Philornis* searching for nestlings. Another behavioral difference might be that males simply invest less in anti-parasitic behaviors like preening and grooming such that they are less likely than females to remove *Philornis*. At present, it is unknown whether male and female Hispaniolan Woodpeckers differ in preening behavior. In at least some bird species, though, males tend to spend more time grooming rather than less (Cotgreave & Clayton 1994, Oswald *et al*. 2019). With respect to morphology, male Hispaniolan Woodpeckers have bills that are on average 25% longer than those of females (unpubl. data; see also Selander 1966), and it could be that their longer bills reduce their effectiveness at removing *Philornis* eggs or larvae. Male Hispaniolan Woodpeckers are also larger in other dimensions of size, including weight, and it could be that their larger size increases the probability that they miss a parasite during preening. Lastly, with regards to immunology, male Hispaniolan Woodpeckers might be more tolerant and/or less resistant to *Philornis* infestation. Widespread evidence exists for sex differences in immunocompetence (e.g., Kelly et al. 2018), but to the best of my knowledge, this possibility remains unstudied with respect to *Philornis*. These alternative behavioral, morphological, and immunological explanations clearly warrant future study.

### Future considerations and implications for *Philornis* biology

Inter-population comparisons of *Philornis* parasitism in Hispaniolan Woodpeckers could be a fruitful course of future research since this woodpecker occupies a range of habitats and elevations. The *Philornis* prevalence on adult woodpeckers documented in the present study was the same as that reported for the species at nearby Rancho Baiguate (H.M. Garrod pers. comm.; Quiroga *et al*. 2020) but higher than that reported from coastal, low elevation Punta Cana (7%; L. Soares and S.C. Latta pers. comm.; Quiroga *et al*. 2020). Whether these differences correspond to *Philornis* population sizes differing according to habitat or climatic conditions could be explored with the Hispaniolan Woodpecker.

Though the difference in nestling and adult *Philornis* infestation intensity is suggestive of a preference for nestlings by botflies, more work is needed to robustly test this hypothesis. In studies of choice and decision-making, confirming the presence of a positive association provides support for a preference hypothesis, but a more discriminating test involves an experimental choice assay (Dougherty 2020). Similar approaches reveal host preferences in insects (e.g., Linn *et al*. 2003). In this case, presenting adult *Philornis* with a choice test between depositing eggs on a nestling or adult bird would be most revealing. Additionally, it is often assumed that parasitizing nestlings yields a higher fitness payoff, but this hypothesis, as far as I know, remains untested.

Given the observed prevalence and intensity of *Philornis* on both nestlings and adults, the Hispaniolan Woodpecker would make an excellent model system to study this parasite’s biology. For example, it would be revealing to conduct *Philornis* exclusion experiments to better understand how non-nesting use of cavities impacts parasitism outside the breeding season or even for non-breeders during the nesting season. The woodpecker’s abundance would also facilitate testing alternative *Philornis* management programs before using them with species of conservation concern.

## Ethics approval

All research activities described here were approved by the Dominican Republic’s Ministerio de Medio Ambiente y Recursos Naturales and conducted in accordance with IACUC protocol 2008-0185 at Cornell University.

## Availability of data and material

The data that support the findings of this study will be openly available in the repository OSF.IO upon acceptance of the manuscript. I provide a temporary link (https://osf.io/unksw/?view_only=4e1cae68cc294fd59a5906a52ee79767) so that editors and reviewers can review the dataset for my manuscript. I will create a publicly registered version of this, including a permanent link and DOI, once the manuscript is accepted.

## Acknowledgments

I gratefully acknowledge the following individuals for providing invaluable assistance in the field: Michelle Angelucci, Haley Boyle, Cecilia Cerrilla, Will Coleman, Aracely Diaz, Lauren Emerson, Neil Gilbert, Amy Janik, Kiera Kauffman, Thomas Lacerda, Alex Lascher-Posner, Mia Larrieu, Kai Larsen, Cedar Mathers-Winn, Kaylee Nelsen, Alyssa Occhialini, Spencer Schubert, Hannah Stapleton, Mitch Walters, Alexa Waterman-Snow, Paris Werner, and Amber Wichtendahl. I thank Walter D. Koenig, Janis L. Dickinson, Paul W. Sherman, Michael S. Webster, H. Kern Reeve, and Irby J. Lovette for providing constructive criticism at various points in the development of the main project that ultimately led to this separate analysis. Breanna L. Bennett, Trey C. Hendrix, Severine B. Hex, Christie Riehl, Amanda G. Savagian, Maria G. Smith, and Alec E. Downie provided much appreciated constructive feedback and discussions that greatly improved the manuscript. I also thank F. Ishtiaq and two anonymous reviewers for their helpful comments on the manuscript. Martín A. Quiroga, Holly M. Garrod, Letícia Soares, and Steven C. Latta provided much appreciated insight into their observations of *Philornis.* I owe much gratitude to Robert Ortíz and the staff at Museo Nacional de Historia Natural in Santo Domingo for assistance with local permitting. The following sources provided funding support: American Ornithological Society Wetmore Award, Cornell Lab of Ornithology Athena Fund, Department of Neurobiology Animal Behavior Research Grant, Society for the Study of Evolution Rosemary Grant Award, and Sigma Xi Grant in Aid of Research. The following fellowships supported me during fieldwork: Charles Walcott Graduate Fellowship, Linda and Samuel Kramer Graduate Student Fellowship, Eleanore Stuart Graduate Fellowship, Andrew ’78 and Margaret Paul Graduate Fellowship, Kramer Graduate Fellowship, Halberstadt Graduate Fellowship, Anne Marie Brown Summer Graduate Fellowship, and Lab of Ornithology Summer Graduate Fellowship.

## SUPPORTING INFORMATION

Additional supporting information may be found in the online version of this article:

**Table S1.** *Individual adult Hispaniolan Woodpeckers recapture records*.

**Table S2.** *Individual adult Hispaniolan Woodpeckers with active* Philornis *infestations*.

**Table S3.** *Sample size breakdown of Hispaniolan Woodpecker adult-nestling* Philornis *concurrent infestations*.

**Figure S1.** *Hispaniolan Woodpecker hatch date histogram with earliest and latest adult capture dates*.

