## SUPPORTING INFORMATION for "Life-stage and sex influence *Philornis* ectoparasitism in a Neotropical woodpecker (*Melanerpes striatus*) with essential male parental care"

### SUPPLEMENTAL METHODS

#### Adult nesting status coding and inclusion/exclusion criteria

In total, I collected 218 adult capture records representing 184 unique individuals (83 females and 101 males). I elaborate here from what is reported in the main text to explain how adults were coded as "nesting" or "not nesting." Adults were coded as "nesting" if they met two criteria: (1) we observed the bird incubating at or provisioning ≥1 nest within the year of capture, *and* (2) we captured the bird after the earliest possible clutch initiation date for its earliest possible nesting attempt within the year of capture. In contrast, the category of birds "not nesting" included (1) birds that never bred within the year of capture, (2) birds captured early in the field season before most nests were initiated (between January and early April), and (3) birds known to have bred in the year of capture but captured before their earliest nesting attempt in that year. Based on these criteria, I assigned nesting status for 193 records. Thus, I omitted 25 records (representing 25 unique individuals) from the original 218 capture records because I lacked adequate data on breeder status. All of the omitted records lacked any direct observation of nesting and included the following: 9 birds for which breeding characteristics (i.e., presence of a brood patch) were not recorded during capture, 5 birds lacking brood patches, 1 bird molting only its abdomen (which could have been re-feathering after development of a brood patch), 6 birds molting the abdomen but also other feather tracts, 1 bird with a smooth abdomen, 2 birds with vascularized brood patches, and 1 bird with a wrinkled brood patch. These last four birds could potentially be considered as "nesting," but I omitted them from the analysis reported to be cautious and only use records with stronger evidence of nesting and/or direct nesting observations.

The remainder of the 193 adult records consisted of 82 observations representing 72 unique females (34 females nesting, 34 females not nesting, and 4 females with observations in both categories) and 111 observations representing 91 unique males (43 males nesting, 43 males not nesting, and 5 males with observations in both categories).

The second analysis included only the subset of adults known to breed within the year of capture and contrasted birds actively nesting with those not nesting at time of capture. This subset included 136 adult records consisting of 61 observations representing 53 unique females (35 females actively nesting, 15 not actively nesting, and 3 with observations in both categories) and 75 observations representing 62 unique males (47 males actively nesting, 14 not actively nesting, and 1 with observations in both categories).

**Table S1.** Individual adult Hispaniolan Woodpecker recapture records. Data presented included Individual's band number (ID), sex (F = female; M = male), capture date, breeder status, and *Philornis* infestation status. Breeder status indicates when an individual was captured relative to any nesting attempts it had during that calendar year.

| ID | Sex | Capture date | Breeder status | *Philornis* infestation status |
|---|---|---|---|---|
| 121 | F | 16 Apr 2014 | before first brood | -No *Philornis* |
| | | 6 Mar 2015 | before first brood | -Old *Philornis* wounds |
| 124 | M | 19 Apr 2014 | before first brood | -No *Philornis* |
| | | 18 Jun 2014 | nestlings | -Old *Philornis* wounds |
| 126 | F | 19 Apr 2014 | nestlings | -No *Philornis* |
| | | 10 Mar 2015 | before first brood | -≥5 old *Philornis* wounds |
| 133 | M | 21 Apr 2014 | before first brood | -No *Philornis* |
| | | 8 Mar 2015 | before first brood | -≥3 old *Philornis* wounds |
| 137 | M | 24 Apr 2014 | before first brood | -No *Philornis* |
| | | 13 Mar 2015 | before first brood | -Old *Philornis* wound |
| | | 7 Apr 2017 | not breeding | -Old *Philornis* wound |
| 231 | M | 21 Jun 2014 | unknown | -No *Philornis* |
| | | 20 Jul 2014 | unknown | -Active *Philornis* wound (2 larvae) and old wounds |
| 241 | M | 26 Jun 2014 | nestlings | -Active *Philornis* wounds (3 larvae) |
| | | 25 Feb 2016 | before first brood | -No *Philornis* |
| 251 | M | 18 Jul 2014 | unknown | -No *Philornis* |
| | | 3 Mar 2015 | before first brood | -Old *Philornis* wounds |
| 269 | M | 25 Mar 2015 | not breeding | -Old *Philornis* wound |
| | | 15 Jun 2015 | not breeding | -No *Philornis* |
| 370 | F | 31 May 2015 | nestlings | -No *Philornis* |
| | | 19 Jul 2015 | nestlings | -Active *Philornis* wound (1 larva) |
| 411 | M | 20 Jun 2015 | nestlings | -No *Philornis* |
| | | 4 Aug 2016 | after nesting | -Old *Philornis* wounds |

**Table S2.** Adult Hispaniolan Woodpeckers with active *Philornis* infestations (i.e., current wounds containing ≥1 larva). Data presented included Individual's band number (ID), sex (F = female; M = male), capture date, and nesting status at the time of capture.

| ID | Sex | Capture date | Nesting status | Additional notes |
|---|---|---|---|---|
| 196 | F | 2-Jun-2014 | Nestlings (~29 d post-hatch) | 2 of 2 nestlings infested: 4 and 7 larvae on each |
| 231 | M | 20-Jul-2014 | Unknown | Observed attacking nestlings on 21-Jun-2014; also captured on that date and had no *Philornis* |
| 241 | M | 26-Jun-2014 | Nestlings (~22 d post-hatch) | 4 of 4 nestlings infected: 9–13 larvae per nestling |
| 263 | F | 8-Mar-2015 | Not breeding | Brood patch absent; had 3 scars and 1 active wound |
| 367 | M | 10-May-2017 | Unknown | Smooth abdomen; dead *Philornis* still in wound |
| 370 | F | 19-Jul-2015 | Nestlings (~11 d post-hatch) | 3 of 3 nestlings infested: 8–9 larvae per nestling |
| 427 | M | 20-Jul-2015 | Nestlings (~11 d post-hatch) | Mate of F 370 |
| 429 | M | 21-Jul-2015 | Nestlings (~13 d post-hatch) | 2 of 2 nestlings infested: 1 and 7 larvae on each; same nest tree as F 370 and M 427 |
| 501 | M | 7-Apr-2016 | Unknown | Brood patch absent; 1 fresh wound lacking larva |
| 633 | M | 25-Jun-2017 | Post-fledging | Nestlings fledged between 23 and 25 May 2017; 1 of 2 nestlings infested with 1 larva |
| 703 | M | 1-Jul-2017 | Unknown | Wrinkled brood patch; no known nest |
| 707 | M | 4-Jul-2017 | Nestlings (~20 d post-hatch) | 4 of 4 nestlings infested: 1–3 larvae per nestling |

**Adult-nestling concurrent infestation details**

Pertaining to the prediction (P2.3) that parents with *Philornis* infested nestlings are more likely to be infested than parents with non-infested nestlings, I analyzed the infestation status of parent birds for 41 nests represented by 40 unique parents or parent pairs. These observations included a mix of nests for which one parent ($n$ = 22 nests, 8 represented by the mother and 14 represented by the father) or both parents ($n$ = 19 nests) were sampled (Table S3). This data set included 58 unique adults, including 25 unique females (2 females were sampled twice: 1 during two different nesting attempts and 1 during the same nesting attempt) and 33 unique males (1 male was sampled twice during the same nesting attempt).

**Table S3.** Sample size breakdown of Hispaniolan Woodpecker adult-nestling *Philornis* concurrent infestation. Parental sampling included either one or both parents. Two counts are presented for each sampling: first, the number of nests with ≥1 adult showing any evidence of *Philornis* (all wounds) and, second, the number of nests with ≥1 adult with current infestations (active wounds). Nestling status includes whether nestlings had any *Philornis* wounds (infested) or none (not infested).

| | Nests observed | | | | | |
|---|---|---|---|---|---|---|
| | One parent sampled | | | Both parents sampled | | |
| **Nestling status** | **All wounds**[*] | **Active wounds**[*] | ***n*** | **All wounds**[*] | **Active wounds**[*] | ***n*** |
| Infested | 5 (1,4) | 3 (1,2) | 10 | 4 (1[**],4[**]) | 2 (1[†],2[†]) | 7 |
| Not infested | 1 (0,1) | 0 | 12 | 3 (1,2) | 0 | 12 |

[*]Numbers in parentheses correspond to the number of adult females and males, respectively.
[**]Two parents from the same nest were infested.
[†]Two parents from the same nest had active wounds.

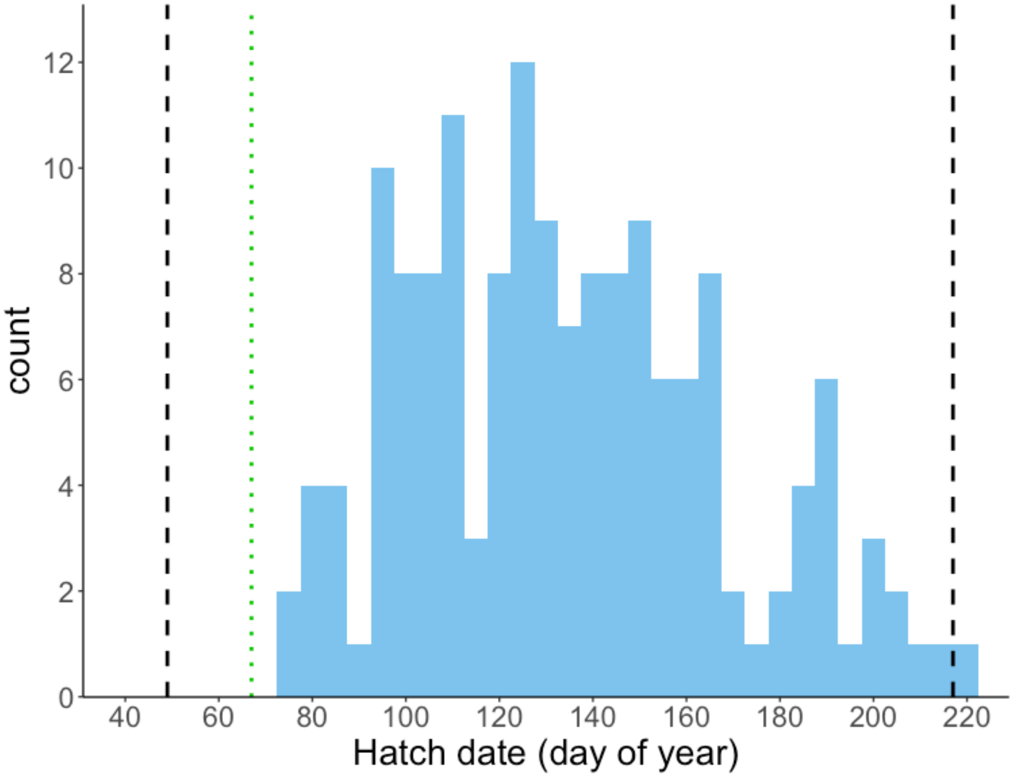


**Figure S1**. Histogram showing hatch date (day of year) for Hispaniolan Woodpecker nests. The black dashed lines represent the earliest and latest capture dates for adult birds included in the present study, and the green dotted line represents the earliest capture date associated with an active *Philornis* infestation on an adult. The mean ± standard error date of capture for adults was 127 ± 3 (i.e., May 7 ± 3 d).
